# Expression of non-neuronal Tau in humans and mice

**DOI:** 10.64898/2026.02.27.708441

**Authors:** Christiana Lekka, Michael Ellis, Katie Holden, Christine S. Flaxman, John A. Todd, M. Irina Stefana, Sarah J. Richardson

## Abstract

**Background:** Tau, encoded by the microtubule-associated protein tau gene (*MAPT*), is traditionally associated with neuronal function in maintaining cytoskeletal stability and regulating intracellular transport. Its dysregulation is implicated in a range of neurodegenerative disorders. Beyond its well-established high levels of expression in neurons of the central and peripheral nervous systems, increasing evidence indicates that Tau is also expressed in non-neuronal cells/tissues such as muscle, kidney, testis, and pancreas, albeit at much lower levels than in neurons. However, the assessment of low levels of non-neuronal Tau requires that the specificities of the anti-Tau antibodies used are fully validated.

**Methods:** Three previously validated anti-Tau antibodies (Tau-12, Tau-1, RD3) were used to test Tau expression levels in 33 mouse and 66 human tissues using Food and Drug Administration-quality tissue microarrays in immunocytochemistry and western blot analyses.

**Results:** The presence of Tau was confirmed in salivary gland, kidney, skeletal muscle, heart, pancreas and oesophagus in both human and mouse tissues.

**Conclusions:** Dysregulation of Tau protein expression and alterations in its post-translational modification is a causative factor in neurodegenerative disease. We have demonstrated a wider Tau expression in other tissues and cells, outside the brain, which may be dysregulated in other diseases.

## 1. Introduction

Tau, encoded by the microtubule-associated protein tau gene (*MAPT*), is traditionally associated with neuronal function in maintaining cytoskeletal stability and regulating intracellular transport. Its dysregulation is implicated in a range of neurodegenerative disorders, including Alzheimer’s disease (AD), Pick’s disease (PiD), Parkinson’s disease (PD) and Progressive Supranuclear Palsy (PSP) (Arendt, Stieler, and Holzer 2016). Beyond its well-established high levels of expression in the central and peripheral neuronal cells (CNS and PNS), increasing evidence indicates that Tau is also expressed in non-neuronal cells in peripheral tissues, such as muscle, kidney, testis, and pancreas, albeit at much lower levels than in neurons (Ashman et al. 1992; Maj et al. 2010; Murakami et al. 1995; Shults et al. 2020; Sigala et al. 2014; Valles-Saiz et al. 2022). These observations raise questions around its possible functional significance in non-neuronal cells and in diseases including cancer, muscle disorders and type 2 diabetes (T2D), which is commonly associated with AD and PD. In addition, owing to the lower expression levels, the specificity and sensitivity of anti-Tau antibodies become a critical feature of investigations into non-neuronal expression. An issue we have reported on previously (Ellis et al. 2024).

The complexity of Tau protein biology is due to its diverse isoforms, arising from alternative splicing of the MAPT gene, and its extensive post-translational modifications (PTMs), such as phosphorylation, acetylation, and truncation. These modifications significantly influence Tau’s localisation, interactions, and functional roles, and further emphasise the need for careful selection of antibodies. Many widely cited PTM-agnostic “total” anti-Tau antibodies exhibit cross-reactivity or sensitivity to PTM states, leading to inconsistent results. Our previous work highlighted these issues and established a robust antibody selection framework—the “traffic light system (TLS)”—to prioritise antibodies based on specificity, sensitivity, and suitability for their intended application, including immunohistochemistry (IHC) and western blotting (WB) (Ellis et al. 2024).

The present study aims to address the gaps in knowledge regarding Tau expression and isoform diversity in non-neuronal cells in peripheral tissues using validated antibodies characterised through our TLS, we have investigated Tau expression using Food and Drug Administration-quality human and mouse tissue microarrays (TMAs) in IHC and WB analyses. This approach enables a more reliable characterisation of total Tau and its isoforms across a wide range of tissues, providing a step forward in understanding Tau biology and disease susceptibility beyond neurons.

## 2. Materials and Methods

### 2.1 Tissue

FFPE frozen TMAs comprising 22 different normal tissue types from a cohort of 66 adult donors were commercially available from AMS Biotechnology Europe Ltd (AMSBIO, catalogue number T8234701-1, Table S1). These were stored under HTA Licence 12276.

FFPE mouse TMAs consisting of 18 normal tissue types sourced from the Institute of Cancer Research (ICR) mice were commercially available from Novus Biologicals (catalogue number NBP2-30225, Table S2). The choice of ICR mice is particularly advantageous due to their extensive genetic diversity, closely resembling that found among human populations. The ICR mice were sacrificed at 8 weeks and tissues were fixed with 4% paraformaldehyde.

Dissected mouse pancreas was formalin-fixed and embedded in paraffin blocks, and 4 μM sagittal sections were cut from a 9-month-old male from a wildtype (WT) C57Bl/6J background. All procedures were approved by local ethical review panels and conducted under UK Home Office project licenses in accordance with the Animals (Scientific Procedures) Act 1986.

### 2.2 Antibodies

A panel of anti-Tau antibodies against total Tau forms (Tau-12 Millipore #MAB2241), dephosphorylated Tau forms (Tau-1 Millipore #MAB3420) and isoform specific Tau forms directed against 3R isoform (RD3 Millipore #05-803) were selected (Table1).

### 2.3 Immunohistochemistry

Human and mouse TMAs were dewaxed, rehydrated, and permeabilised in graded ethanol to 50% ethanol. Heat-induced epitope retrieval (HIER), using a 10 mM citrate buffer (pH6.0), was performed to unmask the epitopes by placing the sections in a plastic pressure cooker in a microwave oven on full power for 20 min. The sections were treated with lambda (λ) phosphatase (λPP) (10,000 U/ml), 1 mM MnCl2 (Biolabs #B17615) and 1x PMP buffer (Biolabs #B07615) overnight, followed by an additional HIER to stop the λPP reaction. The sections were then blocked with 5% normal goat serum (NGS) and/or mouse-on-mouse blocking reagent (#MKB-2213, 1 drop in 1 ml TBS) and incubated with the appropriate primary antibody (Table 1). The sections were then blocked with Agilent Dako REAL peroxidase blocking solution (S202386-2), probed with the secondary antibody Agilent Dako REAL EnVision-HRP, Rabbit/mouse (Agilent Dako, Cat. #K500711-2) and stained with 3,3-Diaminobenzidine (DAB) substrate working solution according to the manufacturer’s instructions. After haematoxylin (Sigma-Aldrich, Cat. #H3136), 2% copper sulphate (CuSO4), Scott’s Tap Water Substitute (STWS) and counterstain, the sections were dehydrated and mounted using Dibutylphthalate Polystyrene Xylene (DPX) (Sigma-Aldrich, Cat. #06522) mounting medium. A human TMA slide was stained solely with the secondary antibody and DAB chromogen, excluding the primary antibody, to discern genuine signals from those possibly attributed to DAB staining. Sections were scanned using the Akoya PhenoImager at X40 magnification.

**Table 1:** Antibodies used in this study.

| Antibody | Supplier | Cat. Number | IHC Conditions | WB conditions |
| --- | --- | --- | --- | --- |
| Tau-12 | Merck Millipore | MAB2241 | 1:500 | N/A |
| Tau-1 | Merck Millipore | MAB3420 | 1:200 | 1:1,000 |
| RD3 | Merck Millipore | 05-803 | 1:500 | N/A |

### 2.4 Image analysis

Whole slide scans were processed using the InForm™ image analysis platform (Akoya Biosciences, US). Quantification analysis of the DAB immunolabelling was performed using the Area Quantification v2.4.9 module from HALO® (Indica Labs), a gold standard image analysis platform for quantitative analysis of IHC data. Statistical significance was determined using one-way ANOVA followed by Tukey’s post hoc multiple comparisons test, with adjusted p-values (p.adj) reported and significance levels annotated as ***p ≤ 0.001, **p ≤ 0.01, *p ≤ 0.05, and ns for p > 0.05.

### 2.5 Western blot

Human tissue lysates were purchased from Zyagen (obtained from certified tissue banks in the USA) and stored at −80°C (Table S3). Lysates were thawed and diluted 3:1 with 4X laemmli buffer (Biorad, cat. no. 1610747) containing β-mercaptoethanol and heated at 95 °C for 10 min. Proteins were separated by sodium dodecyl sulfate– polyacrylamide gel electrophoresis (SDS-PAGE) on 4–15% gradient Mini-PROTEAN TGX Stain Free Gels (Bio-Rad cat. no. 4561086 for 15-well gels) and transferred to Immobilon-FL PVDF membranes (Merck Millipore, cat. no. IPFL00010). To dephosphorylate membranes, membranes were treated with Lambda Protein Phosphatase (λPP, NEB cat no. P0753L) (2000 U enzyme per 1 mL buffer) for 24 hours at RT on an orbital shaker, according to manufacturers instructions. Untreated membranes were mock-treated with equivalent volume of 50% molecular biology-grade glycerol in water. Membranes were blocked in blocking buffer: 5% Amersham™ ECL Prime Blocking Reagent (SLS, cat. no. RPN418) diluted in TBS (no detergent). Primary antibodies were diluted in blocking buffer containing 0.1% Tween-20 and membranes were incubated with primary antibodies overnight at 4 °C. Secondary antibodies were diluted 1:10,000 in blocking buffer containing 0.1% Tween-20 and 0.01% SDS and membranes were incubated for 1 hour at RT. Membranes were imaged dry an Odyssey CLx scanner (LI-COR Biosciences), and blot images were visualised and quantified in the Image Studio software (version 5.2.5; LI-COR Biosciences).

## 3. Results

### 3.1 Antibody selection and criteria for signal assessment

For the investigation of Tau protein expression in peripheral tissues, we selected three antibodies based on criteria established in our previous study’s antibody validation framework: Tau-12, which recognises the N-terminal region of Tau, Tau-1, which detects Tau that is not phosphorylated at residues Ser195 to Thr205, located in the mid-domain of the Tau protein, and RD3, which recognises the 3R Tau isoforms (Table 1).

Given that Tau is a highly phosphorylated protein, and PTM-agnostic ‘total’ antibodies may show variability due to phosphorylation, all TMAs were pre-treated with lambda (λ)phosphatase (λPP) to remove phosphate groups. This treatment aimed to reduce the potential interference caused by phosphorylation, ensuring that the selected antibodies would bind to Tau consistently across both WB and IHC applications.

### 3.2 Tau expression in human and mouse central nervous system

Human TMAs were treated with λPP and immunohistochemical analysis utilizing Tau-12 and Tau-1 antibodies consistently demonstrated positive and comparable immunostaining patterns in the cerebellum, cerebrum and pituitary, as expected (Fig 1A). In alignment with published data (Ellis et al. 2024), the RD3 antibody recognised Tau in the adult human brain (Figure 1A). All three antibodies showed Tau staining in the nucleus of a subset of cells in all three brain regions (Fig 1A, red arrows), in accordance with reports suggesting a nuclear role for Tau (Anton-Fernandez et al. 2023; Younas et al. 2023; Sultan et al. 2011; Maina et al. 2018). The specificity and suitability of the antibodies were further confirmed by detecting Tau in human peripheral nerves (Fig 1A).

**Fig 1.**
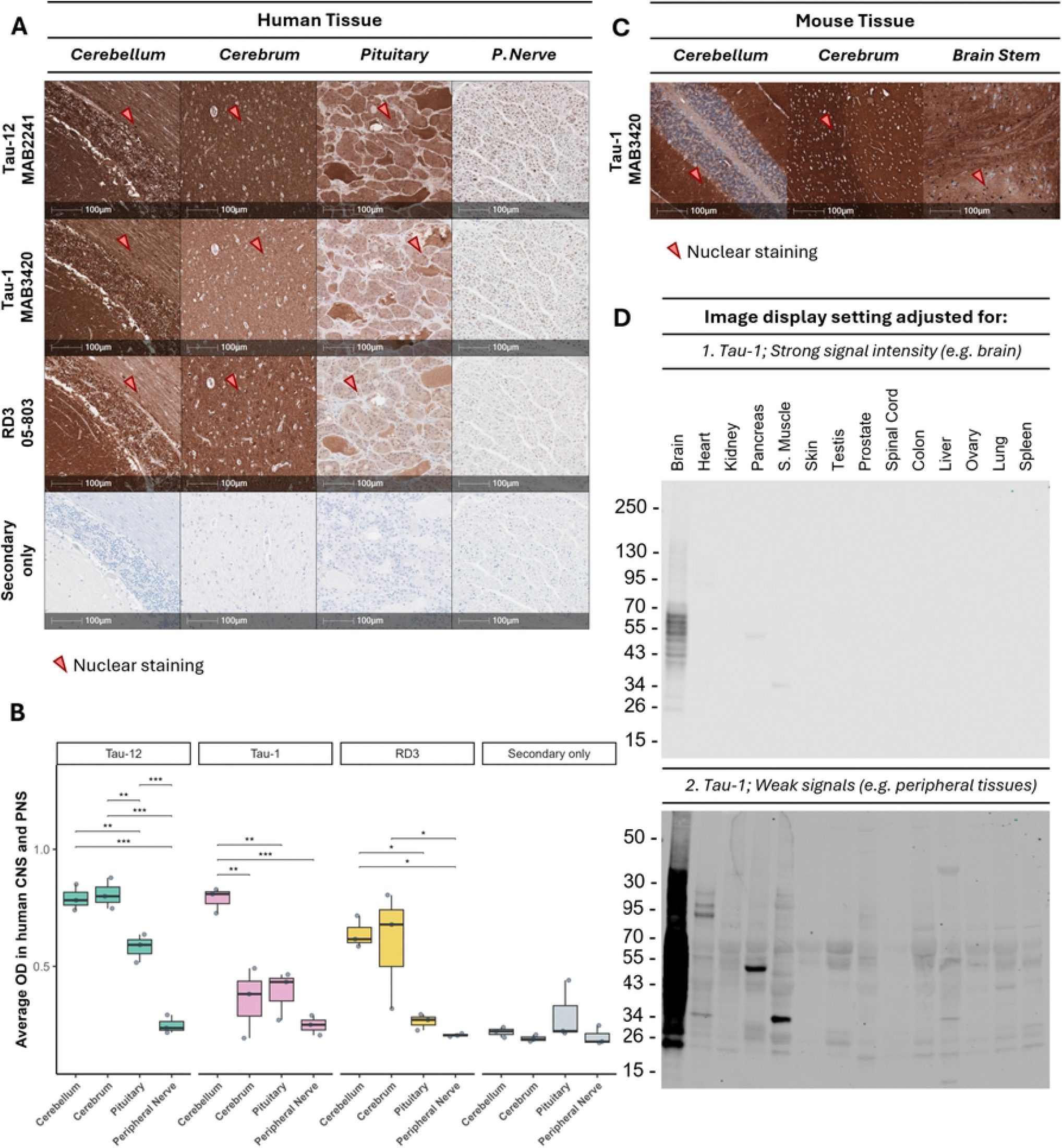
Micrographs demonstrating the cellular localisation of Tau protein forms in the central and peripheral nervous systems. **(A)** Sections of human tissue microarrays (TMAs) were treated with λPP and stained with one of the following antibodies: Tau-12, Tau-1 and RD3; cerebellum, cerebrum, pituitary, and peripheral nerve. A fourth slide was stained solely with the secondary antibody probed with 3,3’-diaminobenzidine (DAB). Whole slide scans were imaged at x40. Scale bars 100 μm. **(B)** Quantification of the optical density of Tau antibodies in the human CSN and PNS. Statistical analysis was performed using the One-way ANOVA with Tukey’s post hoc test. Adjusted p ≤ 0.05 considered significant (p adj ≤ 0.001 ***, ≤ 0.01 **, ≤ 0.05 *, ns). **(C)** Sections of mouse TMAs were treated with λPP and stained with Tau-1; cerebellum, cerebrum, and brain stem. Whole slide scans were imaged at x40. Scale bars 100 μm. **(D)** Immunoblotting using Tau-1 antibody to detect Tau in human tissues.

Tau immunoreactivity in human CNS and PNS was quantified as the average optical density (OD), which is defined as the average intensity of all DAB-positive pixels for each one of the antibodies (Fig 1B). Quantification of OD demonstrated strong immunolabeling in the cerebellum, pituitary, and peripheral nerves with all three antibodies (Fig 1B, Table S4). Tau-12 demonstrated strong immunoreactivity in both the cerebellum and cerebrum at comparable levels, with significantly lower intensity observed in the pituitary gland, followed by the peripheral nerve. Tau-1 showed the highest immunolabeling intensity in the cerebellum, and comparatively weaker staining was detected in the other regions. RD3 exhibited intense immunoreactivity in the cerebellum and cerebrum, whereas staining in the pituitary gland and peripheral nerve was less pronounced, similar to Tau-12. These findings suggest that the mid-region of Tau, recognized by Tau-1, may be masked by PTMs or binding partners in the cerebellum. The 3R Tau isoform, detected by RD3, may be less prevalent in the pituitary. As expected, Tau OD range in the CNS is 0.2-0.9, whereas it was lower in the PNS is 0.2-0.3.

In addition to the human TMAs, mouse TMAs were also treated with λPP and stained with Tau-1 antibody, which identified mouse Tau in the cerebellum, cerebrum and brain stem of ICR mice, as expected (Fig 1C). Nuclear staining with Tau-1 is also observed in a subset of cells in all mouse sections (Fig 1C, red arrows). Tau OD in mouse CNS was quantified; however, statistical analysis was not performed due to limited sample size. As expected from our previous work (Ellis et al. 2024), all antibodies detected Tau in the human CNS and PNS and the mouse CNS, with the secondary-only antibody control in the TMA being negative in all tissues.

To explore the relative expression levels between the brain and various peripheral tissues, human protein lysates were treated with λPP and stained using Tau-1 (Fig 1D). The unmodified full-length Tau has a molecular weight (MW) of 50 to 70 kDa. Strong intensity bands were detected spanning between 50-70kDa in the brain, confirming the specificity of Tau-1 (Fig 1D, top panel). With increased exposure times, strong bands spanning the 50-70 kDa range were detected in the human pancreas and weaker ones in the heart, kidney, skeletal muscle, testis, prostate, colon, liver, ovary, lung and spleen (Fig 1D, bottom panel). Low-MW Tau variants migrating in the 26-40 kDa apparent MW range were detected in the heart, skeletal muscle and liver. Collectively, these data suggest that Tau is prevalent in the brain, and its presence in peripheral tissue is lower relative to the brain. These findings raise questions about whether Tau is expressed in these target tissues in subsets of cells or whether the signal arises due to dense innervation.

### 3.3 Tau expression in human and mouse peripheral tissues

To explore the expression of Tau in human and mouse peripheral tissues, TMAs of normal human and mouse organs were pre-incubated with λPP and immunostained with Tau antibodies. Human TMAs were immunostained with Tau-12, Tau-1 and RD3 (Fig 2A), whereas mouse TMAs were immunostained with Tau-1 (Fig 2B). Tau immunoreactivity was detected within nerves in both species as expected when present (blue arrows). Importantly, in both species, Tau was also present in subsets of non-neuronal cells in the salivary gland, kidney, skeletal muscle, and heart. Tau was present in the human pancreatic islets and oesophagus but was not visually present in the corresponding mouse tissues. However, in the mouse pancreas, no islets were present within the core (Fig 2B). To address this and to confirm whether Tau is present in mouse islets, λPP-treated wild-type (WT) mouse pancreas sections were immunolabelled with Tau-1 (Fig 2C), clearly demonstrating the presence of Tau within mouse islets in agreement with earlier studies (Benderradji et al. 2022; Wijesekara et al. 2018).

**Fig 2.**
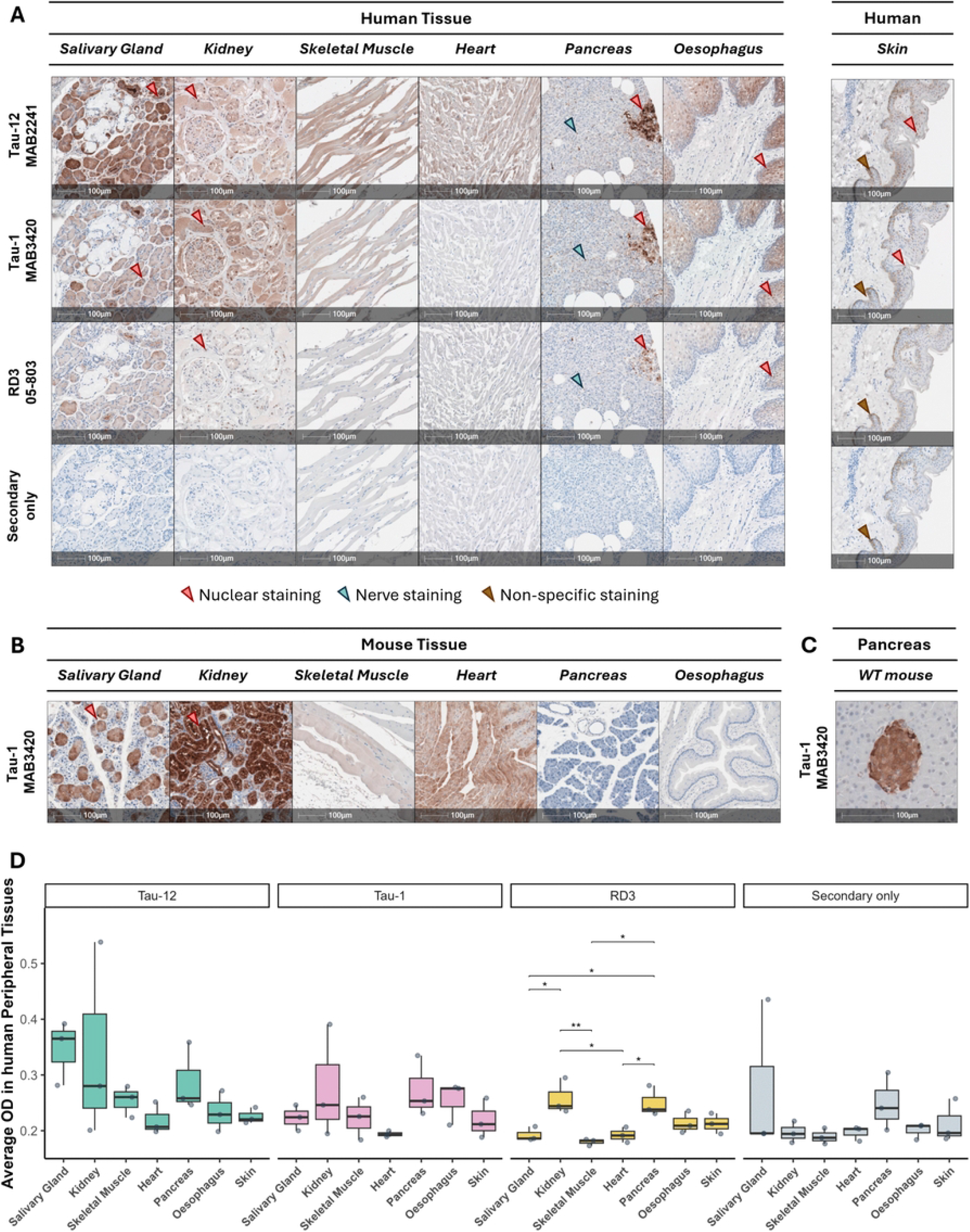
Micrographs demonstrating the cellular localisation of Tau protein forms in the central and peripheral nervous systems. **(A)** Sections of human tissue microarrays (TMAs) were treated with λPP and stained with one of the following antibodies: Tau-12, Tau-1 and RD3; cerebellum, cerebrum, pituitary, and peripheral nerve. A fourth slide was stained solely with the secondary antibody probed with 3,3’-diaminobenzidine (DAB). **(B)** Sections of mouse TMAs were treated with λPP and stained with Tau-1; cerebellum, cerebrum, and brain stem. **(D)** Wild-type mouse pancreas was treated with λPP and stained with Tau-1. Whole slide scans were imaged at x40. Scale bars 100 μm. **(D)** Quantification of the optical density of Tau antibodies in the human CSN and PNS. Statistical analysis was performed using the One-way ANOVA with Tukey’s post hoc test. Adjusted p ≤ 0.05 considered significant (p adj ≤ 0.001 ***, ≤ 0.01 **, ≤ 0.05 *, ns).

Tau expression is predominantly localised to the cytoplasm of most Tau-positive cells. Nuclear localisation of Tau was, however, observed in the salivary gland, kidney and pancreas in both species and in the human oesophagus. In all instances except for the human skin TMAs, the secondary-only slide consistently remained devoid of staining. The staining observed in skin epithelial cells in the secondary-only slide was comparable across all antibodies and, as such, was considered non-specific (Fig 2A). As in the brain (Fig 1A), nuclear Tau immunostaining was evident in a subset of cells with all antibodies but was absent in the secondary-only control slide. This observation indicates the potential expression of Tau within the nucleus of epithelial cells but will need to be confirmed by future studies.

The average OD of Tau immunoreactivity was mostly comparable across human peripheral tissues (Fig 2D, Table S5). Tau-12 and Tau-1 immunoreactivity remained comparable across all tissues. Notably, RD3 immunoreactivity exhibited significant differences between several tissue pairs with kidney and pancreas demonstrating the strongest signal, followed by the kidney and the oesophagus. The weakest signal was reported in the salivary gland, the skeletal muscle and the heart. These findings suggest an overall uniform Tau expression across these peripheral tissues, proposing that the distribution of Tau could be isoform-specific.

Tau immunoreactivity appeared to be restricted to the nerves in the prostate, stomach, thymus, colon, ovary, small intestines and liver (Fig 3). No positive signal was observed in the testis, spleen, lung and bone marrow (Fig 4), contrary to published literature supporting Tau expression in the human testis and lung (Shults et al. 2020; Sigala et al. 2014).

**Fig 3.**
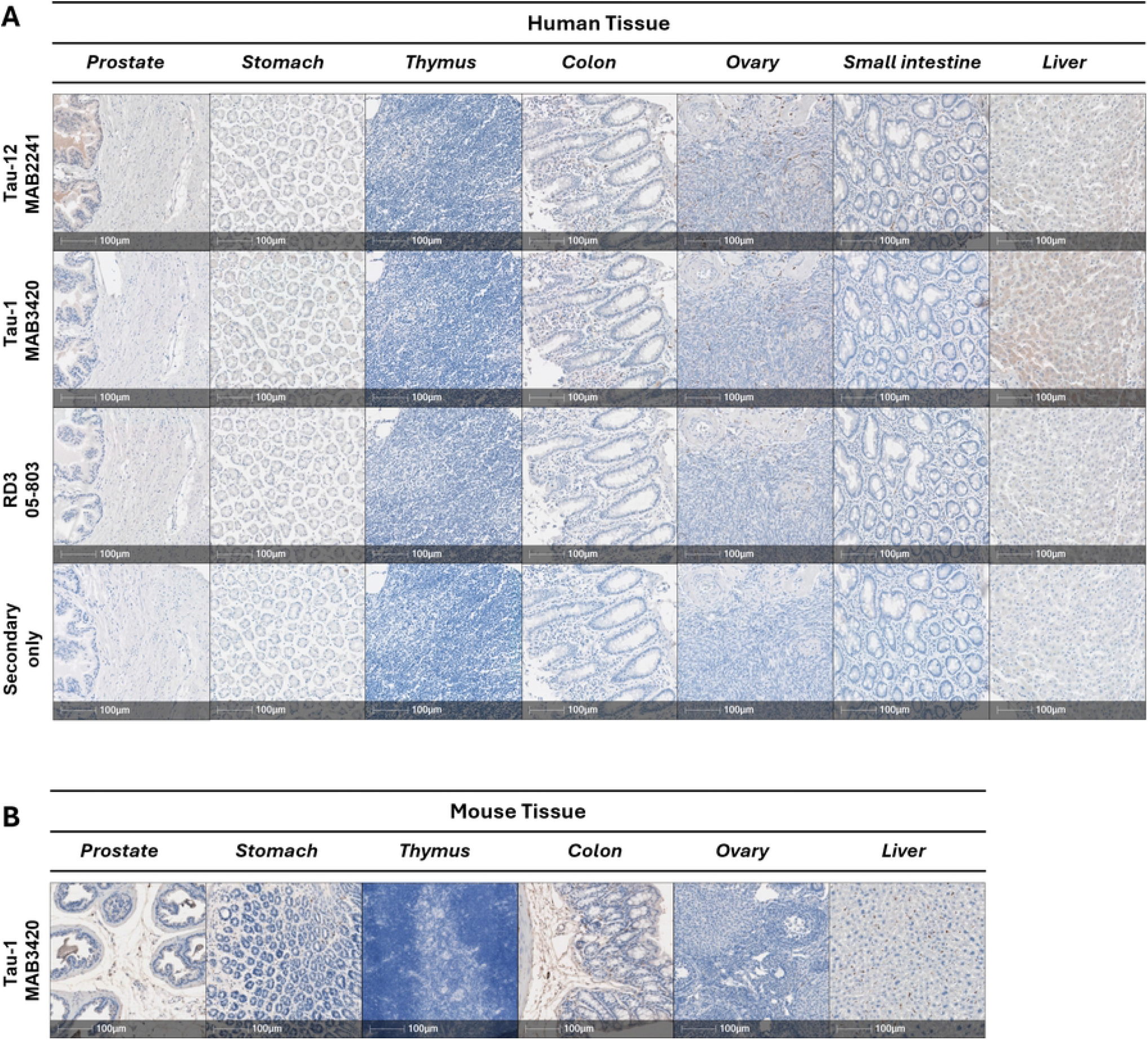
Micrographs demonstrating the cellular localisation of Tau protein forms in the central and peripheral nervous systems. **(A)** Sections of human tissue microarrays (TMAs) were treated with λPP and stained with one of the following antibodies: Tau-12, Tau-1 and RD3; prostate, stomach, thymus, colon, ovary, small intestine and liver. A fourth slide was stained solely with the secondary antibody probed with 3,3’-diaminobenzidine (DAB). **(B)** Sections of mouse TMAs were treated with λPP and stained with Tau-1; prostate, stomach, thymus, colon, ovary and liver. Whole slide scans were imaged at x40. Scale bars 100 μm.

**Fig 4.**
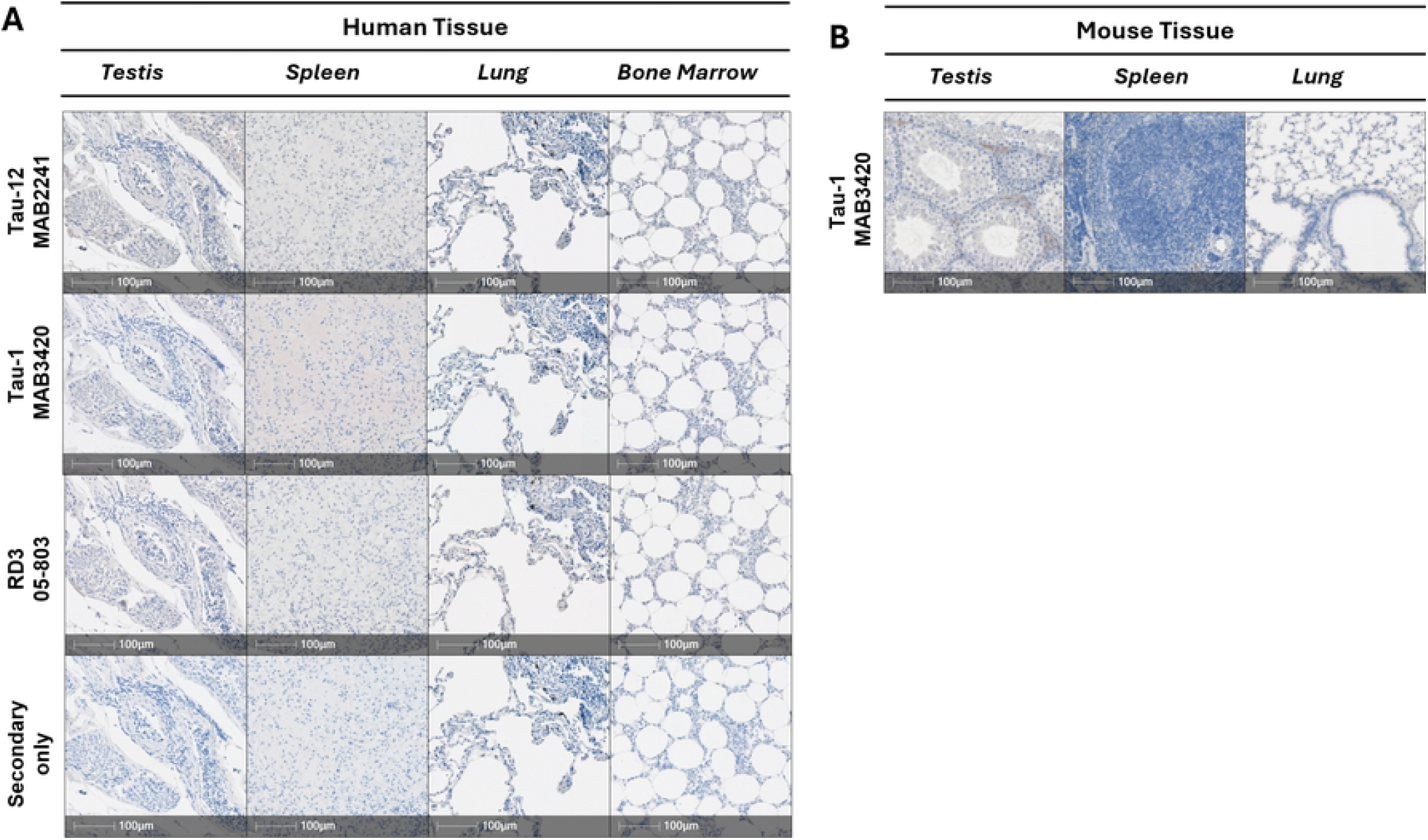
Micrographs demonstrating the cellular localisation of Tau protein forms in the central and peripheral nervous systems. **(A)** Sections of human tissue microarrays (TMAs) were treated with λPP and stained with one of the following antibodies: Tau-12, Tau-1 and RD3; testis, spleen, lung, and bone marrow. A fourth slide was stained solely with the secondary antibody probed with 3,3’-diaminobenzidine (DAB). **(B)** Sections of mouse TMAs were treated with λPP and stained with Tau-1; testis, spleen, and lung. Whole slide scans were imaged at x40. Scale bars 100 μm.

## 4. Discussion

Tau’s function is established as a microtubule-stabilising protein in neurons (Arendt, Stieler, and Holzer 2016). However, emerging data suggest that Tau may contribute to non-neuronal cellular processes, possibly with functions beyond stabilizing axonal microtubules (Maina et al. 2018). Tau has also been observed in the nucleus of non-neural cells, although its nuclear function remains poorly understood (Maina et al. 2018). This raises important questions about its role in cellular functions outside the CNS and PNS. However, the presence of Tau outside the brain has not been systematically studied using well-validated Tau reagents.

Our study provides evidence of Tau expression in peripheral human and murine tissues, including the salivary gland, kidney, skeletal muscle, heart, pancreas, and oesophagus. WB analysis reveals that Tau is expressed at lower levels in human peripheral organs compared to the brain, underscoring the need for highly sensitive antibodies that can reliably detect low endogenous Tau levels. To further determine whether the weak peripheral staining arises from Tau expression within specific cell types or from dense innervation, we employed immunostaining to enable localisation of Tau staining within the different tissues. This approach confirmed the presence of Tau in multiple human and murine peripheral organs, including the salivary gland, kidney, skeletal muscle, heart, pancreas, and oesophagus. These findings suggest that Tau expression in peripheral tissues may be cell-type specific rather than species-specific.

We note the expression of Tau in mouse and human pancreas. Tau’s role in insulin regulation in the pancreas has gained attention, especially in the context of β-cell function (Ho et al. 2020; Maj et al. 2010; Mangiafico et al. 2023). The expression of Tau in murine β cells has been reported to regulate microtubule stability and influence glucose-stimulated insulin secretion, underscoring its importance in metabolic processes (Ho et al. 2020). Elevated Tau levels and hyperphosphorylated forms have been linked to insulinomas, indicating a potential role in human insulin secretion and glucose metabolism (Maj et al. 2010). Tau’s involvement in β-cell dynamics and its impact on insulin vesicle trafficking highlight the need for further study into its role in the islet in both health and disease. It is conceivable that Tau could present a novel therapeutic target for disorders such as T2D (Mangiafico et al. 2023; Miklossy et al. 2010) and other metabolic diseases.

In addition to exploring Tau’s distribution, we aimed to address the challenges associated with its detection in peripheral tissues by systematically validating anti-Tau antibodies in both WB and IHC. Our findings indicate that no single “PTM-agnostic” Tau antibody can detect all Tau isoforms. Notably, Tau-12 and Tau-1 exhibited comparable staining patterns across different brain regions but differed in their ability to detect Tau in the cerebrum. In peripheral tissues, both antibodies produced a similar staining pattern, suggesting that Tau expression in the cerebrum may be influenced by PTMs beyond phosphorylation or through interactions with specific binding partners. Given the complexity and variability of Tau expression, future studies should prioritise developing and validating antibodies that can detect the full spectrum of Tau isoforms, particularly in peripheral tissues. Additionally, using advanced methodologies, including comprehensive tissue sampling and standardised IHC protocols, will be crucial to overcoming current limitations in TMA analyses and improving the accuracy of Tau detection across different tissues.

In conclusion, our results support a role for Tau functions in non-neuronal cells in peripheral tissues. While our findings provide insights into Tau’s roles outside the CNS and PNS, they also highlight the challenges in fully understanding Tau expression due to its complexity, and the variability of reagent performance, both between species and the recognition of differently modified forms. Future studies should focus on expanding the tissue pool and employing advanced techniques to gain a deeper understanding of Tau’s diverse roles in both health and disease.

## Limitations

This study has several limitations. Firstly, the use of TMAs, which represent only a fraction of the sub-tissue structures, may have restricted our ability to identify all Tau-positive or Tau-negative structures. Expanding the tissue pool and increasing the sample size would enhance the generalisability of our findings. Secondly, by pre-treating with λPP, we are examining the presence/ absence of Tau forms; as such, these findings do not provide information about PTM-forms of Tau that may be differentially present in peripheral tissues. Future studies will be required to decipher this, but crucially, will require careful use of validated antibodies that recognise genuine PTM forms and where the species differences are considered. Lastly, our reliance on normal adult TMAs will not fully capture age- or disease-related variations in Tau isoform expression.

## Acknowledgments

Thank you to Dr Marie-Louise Zeissler and Dr Shalinee Dhayal for earlier work optimising some of the reagents in this study. 2021 EASD-Novo Nordisk Foundation Diabetes Prize for Excellence to JAT.

## Supporting information

**S1 Table. Human tissue microarrays**. Human TMAs – donor ID, organ, age, sex.

**S2 Table. Mouse tissue microarrays**. Mouse TMAs – TMA number, sex, and organ.

**S3 Table. Human lysates**.

**S4 Table. Statistical analysis of optical density in human central and peripheral nervous system**. Differences between groups were analysed using one-way ANOVA followed by Tukey’s post hoc multiple comparisons test.

**S5 Table. Statistical analysis of optical density in human peripheral organs**. Differences between groups were analysed using one-way ANOVA followed by Tukey’s post hoc multiple comparisons test.

## Notes

### Competing Interest Statement

The authors have declared no competing interest.

